# Virus diversity and activity is driven by snowmelt and host dynamics in a high-altitude watershed soil ecosystem

**DOI:** 10.1101/2023.03.06.531389

**Authors:** Clement Coclet, Patrick O. Sorensen, Ulas Karaoz, Shi Wang, Eoin L. Brodie, Emiley A. Eloe-Fadrosh, Simon Roux

## Abstract

Viruses, including phages, impact nearly all organisms on Earth, including microbial communities and their associated biogeochemical processes. In soils, highly diverse viral communities have been identified, with a global distribution seemingly driven by multiple biotic and abiotic factors, especially soil temperature and moisture. However, our current understanding of the stability of soil viral communities across time, and their response to strong seasonal change in environmental parameters remains limited. Here, we investigated the diversity and activity of environmental DNA and RNA viruses, including phages, across dynamics seasonal changes in a snow-dominated mountainous watershed by examining paired metagenomes and metatranscriptomes. We identified a large number of DNA and RNA viruses taxonomically divergent from existing environmental viruses, including a significant proportion of RNA viruses target fungal hosts and a large and unsuspected diversity of positive single-stranded RNA phages (*Leviviricetes*), highlighting the under-characterization of the global soil virosphere. Among these, we were able to distinguish subsets of active phages which changed across seasons, consistent with a “seed-bank” viral community structure in which new phage activity, for example replication and host lysis, is sequentially triggered by changes in environmental conditions. Zooming in at the population level, we further identified virus-host dynamics matching two existing ecological models: “Kill-The-Winner” which proposes that lytic phages are actively infecting abundant bacteria, and “Piggyback-The-Persistent” which argues that when the host is growing slowly it is more beneficial to remain in a lysogenic state. The former was associated with summer months of high and rapid microbial activity, and the latter to winter months of limited and slow host growth. Taken together, these results suggest that the high diversity of viruses in soils is likely associated with a broad range of host interaction types each adapted to specific host ecological strategies and environmental conditions. Moving forward, while as our understanding of how environmental and host factors drive viral activity in soil ecosystems progresses, integrating these viral impacts in complex natural microbiome models will be key to accurately predict ecosystem biogeochemistry.

## INTRODUCTION

Soil microbiomes represent a major reservoir of microbial diversity on Earth, and provide many critical ecosystem services such as driving major the major transformation of carbon, and other nutrients and sustaining plant growth^1^. Soil ecosystems, found across a large range of environments including deserts, wetlands, forests, and mountains, are vulnerable to climate change^2–5^. Mountainous ecosystems in particular are impacted by unprecedented snowpack reductions and earlier spring snowmelt, which can trigger rapid collapse of the microbial biomass and abrupt changes in the composition and functioning of soil microbiomes^6^. However, predicting the impact of environmental changes on soil microbiomes remains challenging^2, 3, 5^, and holistic studies, including soil virus-microbes interactions, are needed to elucidate the ecological consequences of climate change on these ecosystems.

Viruses are commonly found in every biome^7^, from the human gut to the ocean, and in many different soil types including agricultural soils^8–10^, grasslands^11–16^, and deserts^17–19^. The complexity of habitats and heterogeneity of microorganisms in soils, spanning protozoa, algae, fungi, bacteria, and archaea, provide an environment where viruses are highly diverse and may play essential roles in microbe functions^20–22^. Viruses infecting bacteria and archaea (hereafter referred to as phages) are the most common and diverse group of viruses identified in soil^19–22^, and can harbor various virion morphologies and genome types including double-stranded DNA (dsDNA), single-stranded DNA (ssDNA), single-stranded RNA (ssRNA), and double-stranded RNA (dsRNA) genomes^25^. Soil viruses are highly abundant and diverse, however current public databases still capture only a fraction of this diversity (mainly dsDNA viruses)^7, 27–29^, due to a combination of biological biases and methodological limitations. Virus genome representation in public database is also biased towards dsDNA phages, which represent the overwhelming majority of viruses reported in soil to date^21, 22^, while our current knowledge of the diversity and ecological role of soil RNA and ssDNA viruses remains limited^12, 13, 30–32^.

Previous soil viral ecology studies based on viral particle counts and/or multi-omics analyses have suggested that soil warming, permafrost thaw, and shifts in soil moisture directly and/or indirectly influenced soil viral diversity^12, 33, 34^. Overall, spatiotemporal patterns in soil viral community composition could be associated with abiotic (soil temperature and depth, pH, and moisture)^8, 12, 29, 31, 34–39^ and biotic factors such as host community composition^35, 40, 41^. Viruses may impact soil ecosystem functioning especially through viral infections of key biogeochemical-cycling microbes, and these viral-host dynamics may change with environmental conditions^35–37^. Additionally, growing evidence of viral populations carrying auxiliary metabolic genes, i.e. viral-encoded metabolic genes that could provide a fitness advantage to their hosts, provide a complementary way by which viruses likely influence soil biogeochemical cycling^12, 34^.

While these studies of soil viral ecology have highlighted the potential influence of viruses on host community structure, nutrient cycling, and other ecosystem processes, major gaps remain in our understanding of soil viral ecology, especially in mountainous environments where abrupt seasonal changes occur. In particular, the global diversity of phages in mountainous soils, especially of ssDNA and RNA phages, remains poorly constrained, and the activity and dynamics of phages across seasons is poorly understood. Additionally, the ecological models that describe phage-bacteria interactions, i.e., lytic predation according to classical predator-prey Kill-the-Winner (KTW) dynamics and temperate infection according to Piggyback-the-Winner (PTW) appear to be conflicting hypotheses as they both occur in same types of environments but have opposite results, and the mechanisms that trigger the lytic-lysogenic switch remain mostly unknown^42, 43^. Thus, the direct and indirect impacts of climate change on mountainous soil viruses, and the subsequent repercussions on soil microbiome and metabolic processes, are currently unknown.

Here, we aimed to address how early snowmelt and, more globally, seasonal disturbances might influence the diversity, composition, and activity of mountainous soil viromes. To this end, we leveraged existing metagenomic and metranscriptomic data from soil samples collected over a year-long time-course in high-altitude mountainous soils of the East River Watershed (ERW), Colorado, United States^44^. This study site was specifically developed as a testbed to develop coordinated and multiscale research that integrate hydrological, biogeochemical, and microbiological studies, and includes previous investigation of resident bacterial, archaeal, and fungi communities^6^. Here, we further explored these soil microbiomes by identifying virus sequences across metagenomes and metatranscriptomes, and connecting the diversity and activity of both DNA and RNA viruses to changes in major biotic and abiotic parameters. Taken together, these analyses shed light on the contrasted dynamics and potentially different infection strategies of different groups of viruses across seasons, paving the way towards a better understanding of virus impacts on soil microbiomes and a robust integration of virus types and strategies into ecosystem biogeochemical models.

## RESULTS

### High diversity of DNA and RNA phages in the East River Watershed soil dataset

To characterize the soil viral diversity in the East River Watershed (ERW), we analyzed 47 and 43 paired metagenomes and metatranscriptomes, respectively, obtained from samples collected in hillslope locations at three depths over four dates from March 2017 to June 2017 (**Figure 1A, and Supplementary Table 1 and 2**). Using established protocols combining virus sequence detection with VirSorter2^42^, refinement with CheckV^43^, and clustering into non-redundant vOTU^44^, we recovered 4,047 and 5,032 non-redundant DNA and RNA viral genomes, respectively (**Supplementary Table 3 and 4**). Similar to previous soil viromics studies, representative contigs from DNA vOTU included a mix of (near-)complete genomes and genome fragments, with nearly 23% (907) longer than 10 kbp, and 101 identified as high-quality (>90% complete) (**Figure 1B, 1C**). Meanwhile, because RNA virus genomes are typically shorter, a larger proportion (n=1,870, 37%) were identified as high-quality (>90% complete) genomes (**Figure 1B, 1C and 1D**).

**Figure 1.**
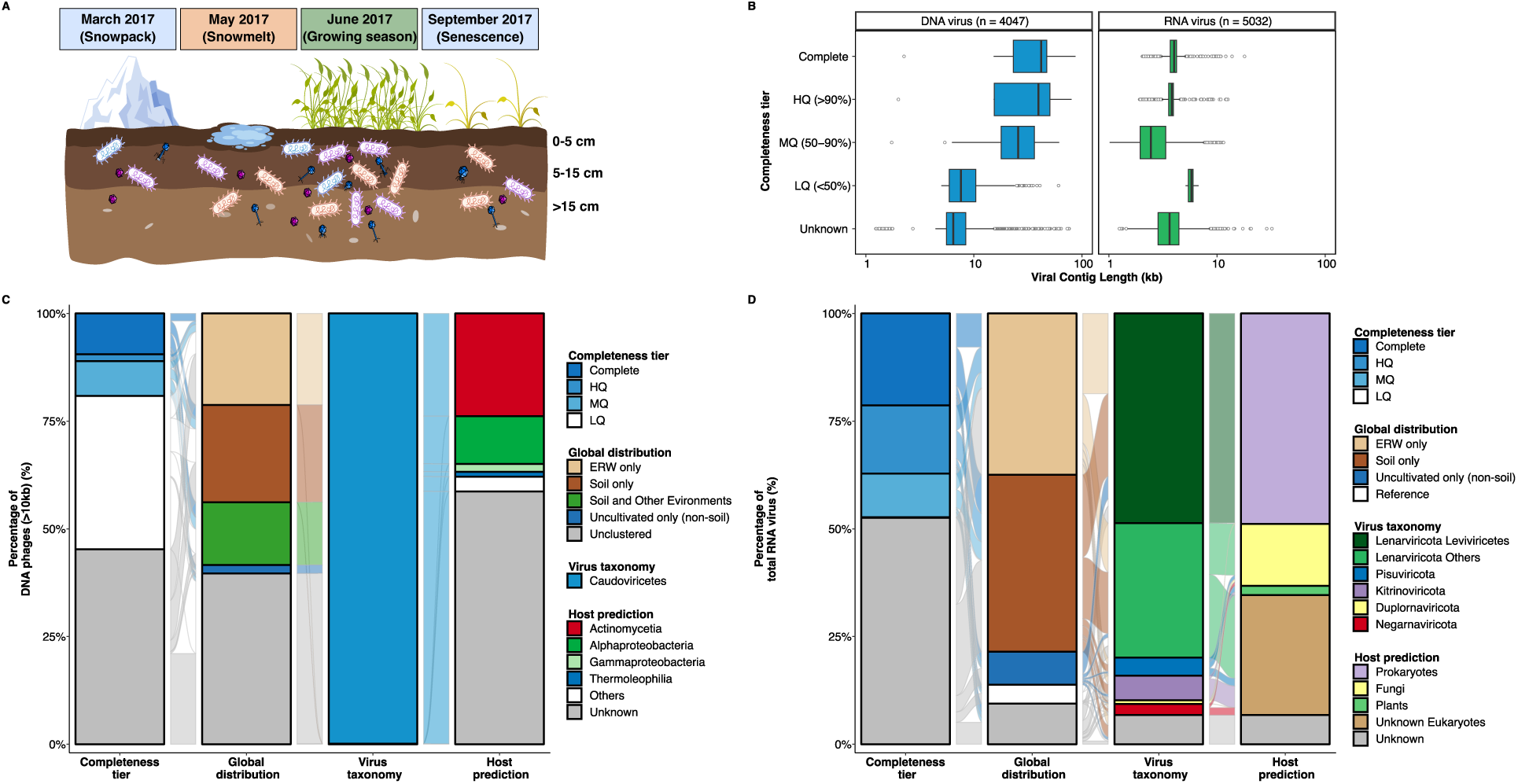
Overview of sampling strategy, and general features of East River Watershed (ERW) DNA and RNA viruses. **A**.Visual schematic of the sampled mountainous ERW soils. Samples were collected from bulk soils in the East River Watershed, Colorado on four dates, starting first at peak winter snow depth (March 7, 2017), during the snowmelt period (May 9, 2017), following the complete loss of snow and the start of the plant growing season (June 9, 2017), and finishing in autumn after plant senescence (September 15, 2017). Soil cores from discrete locations were subsampled and split into three discrete depth increments; 0 to 5 cm, 5 to 15 cm, and 15 cm + below the soil surface. Sample distribution. During snow-free times of the year (i.e., September and June), soils were collected at the upslope and downslope while during periods of winter snow cover (i.e., March and May), snow-pits were dug down to the soil surface in order to sample soils from beneath the snowpack, generating 46 and 43 metagenomic and metatranscriptomic samples, respectively. More information on individual samples is available on ESS-DIVE (https://ess-dive.lbl.gov) using accession numbers listed in Supplementary Table. **B.** Length distribution of the different quality tiers of DNA (blue) and RNA (green) vOTUs, based on their estimated completeness assessed by CheckV. HQ: High-quality (≥90% complete), MQ: Medium-quality (≥50% complete). LQ: Low-quality (<50% complete). **C.** Completeness, global distribution, virus taxonomy, and host taxonomy for DNA vOTUs ≥10kb. Global distribution is based on shared clusters from a vContact2 analysis, virus and host taxonomy are based on GeNomad and iPHoP tools, respectively (see Methods). **D.** Completeness, global distribution, virus taxonomy, and host taxonomy for RNA vOTUs. Global distribution is assessed by genome-based clustering, virus and host taxonomy are based on GeNomad and iPHoP and refined using phylogenetic analyses of the RNA virus marker gene RdRP (RNA-dependent RNA polymerase, see Methods).

A marker gene taxonomic classification performed using GeNomad^48^ suggested that the vast majority (> 90%) of classified DNA vOTUs were bacteriophages from the *Caudoviricetes* class, i.e., tailed dsDNA bacteriophages typically identified in soil metagenomes (**Figure 1C and Supplementary Table 3**). Meanwhile, ∼ 50% of RNA vOTUs were classified into the *Leviviricetes* bacteriophage class, while the remaining 44.6% of vOTUs were assigned across all five recognized phyla of RNA viruses (i.e., *Lenarviricota* outside of the *Leviviricetes* class, *Pisuviricota, Kitrinoviricota, Duplornaviricota,* and *Negarnaviricota*) (**Figure 1D and Supplementary Table 4**). About 5% of RNA vOTUs were not assigned to known RNA virus phyla. To refine this marker-based affiliation of RNA viruses, we performed a phylogenetic analysis of RNA viruses based on the RdRP marker gene (RNA-dependent RNA polymerase)^12, 29, 30, 45^. After clustering ERW RdRP sequences with reference and previously published datasets obtained from soil metatranscriptomes at 50% average amino acid identity (AAI), each cluster representative was assigned to a phylum, and a phylogeny was built for each phylum. Consistent with the marker gene geNomad classification, most of the ERW RdRP grouped within the positive-sense single-stranded *Lenarviricota* (75%), followed by the *Kitrinoviricota* (6.7%) and *Pisuviricota* (5.6%) (**Figure 2A**). The phylogenetic tree of phylum *Lenarviricota* could be further divided into four subclades; the first one corresponds to sequences branching within the *Leviviricetes* class (35.8%), the second and third to sequences branching next to the *Ourlivirales* (11.6%) and *Cryppavirales* (8.8%) orders, while the fourth group represented novel clade with no closely related sequence within the *Wolframvirales* (4.5%) order (**Figure 2B and Supplementary Table 5**). Among the other phyla, *Picornavirales* (2.5%)*, Tolivirales* (2.3%) and *Martellivirales* (1.3%) were the most represented orders (**Supplementary Figure 1A to 1D and Supplementary Table 6 to 9**). Taken together, these results highlight the high diversity of both DNA and RNA viral communities in ERW, with an unsuspected high richness of RNA phages in ERW soils.

**Figure 2.**
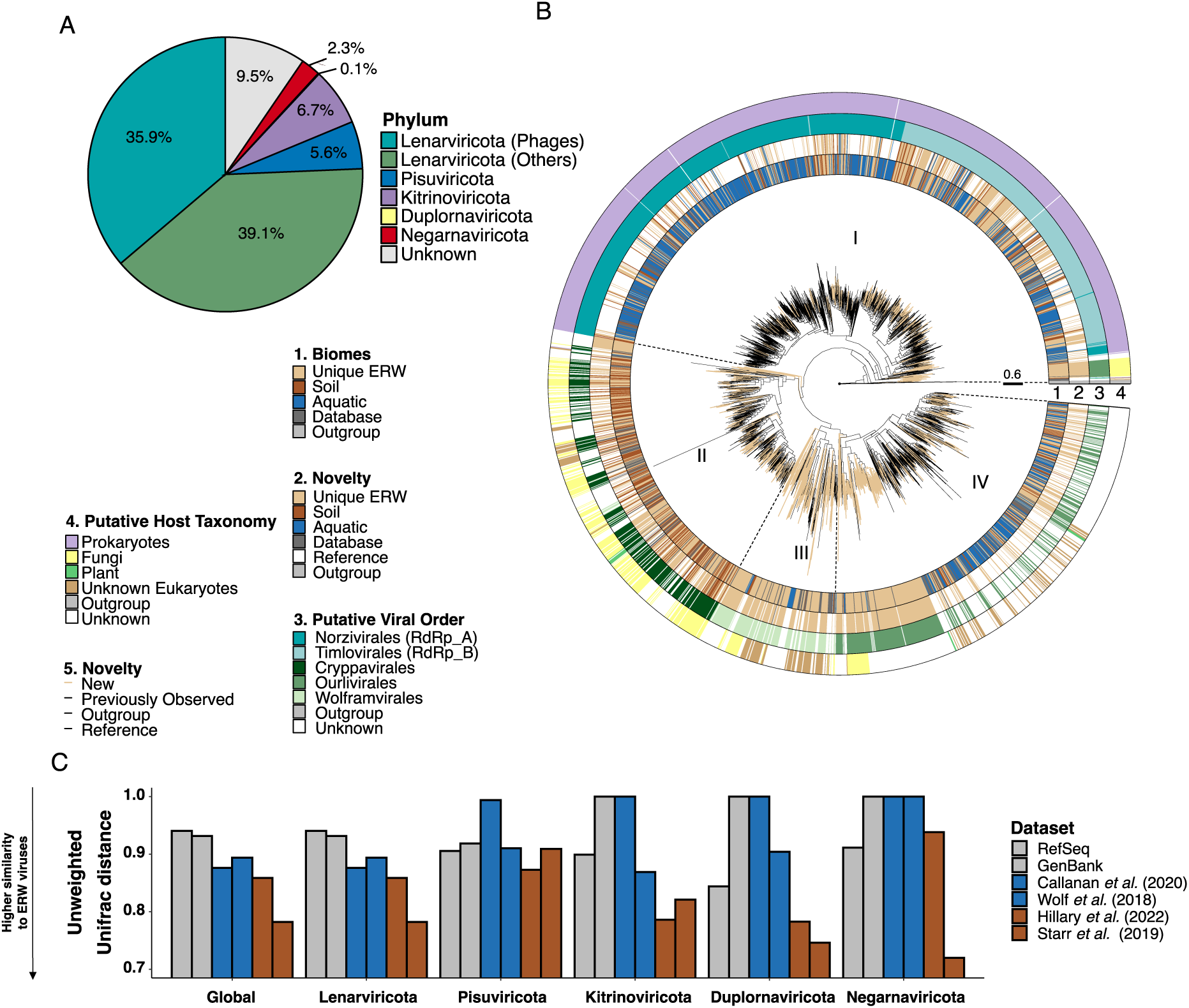
Diversity and phylogenetic analyses of ERW RNA viral communities. **A**. Distribution of RNA viral phyla, based on putative taxonomic assignments for RdRP sequences (derived from viral taxonomy of known RdRP sequences in RefSeq). The *Lenarviricota* phylum is divided between bacteria-infecting viruses (”phages”) and eukaryote-infecting viruses. **B.** Rooted phylogenetic tree of RdRP sequences belonging to the ssRNA *Lenarvirricota* phylum. The tree is rooted using reverse transcriptases as an outgroup and visualized with *ggtree*. The tree contains 1,331 cluster representatives from ERW soil samples (ring 1, light brown), aligned with those used to construct the RNA global phylogeny from previous metatranscriptomic studies and public databases. Clusters composed exclusively of ERW sequences are colored in brown (ring 2) with branches leading to these clusters highlighted in light brown in the tree, while clusters composed of ERW sequences and existing virus sequences are colored by the environment type of the study (soil: dark brown, aquatic: blue, public databases: dark grey). Virus taxonomy (ring 3) and host (ring 4) are predicted based on the position of reference sequences from the RefSeq database in the tree (see Methods). **C.** Unweighted UNIFRAC distances between RdRP sequences identified in this study and previously published collections of environmental RNA viruses^13, 24, 31, 49^. UNIFRAC distances were calculated and are presented separately for each of the 5 RNA virus phyla, with the average distance presented in the first “Global” column. Reference databases are colored in grey, studies from aquatic environments in blue and soils in dark brown. A distance close to 0 means that the two datasets are phylogenetically similar.

### Ecological distribution of ERW soil virus diversity and host connections

We next compared this ERW viral diversity to reference viruses and soil virus metagenomic datasets using either genome-based (vContact2^46^, for dsDNA phages >= 10kb) or gene-based clustering (RdRP, 50% AAI, for RNA viruses). While these analyses included > 12,000 soil DNA phages and 613,000 RNA virus sequences, 20% of the ERW dsDNA phages (**Supplementary Table 10**) and 37% of the ERW RNA viruses were found in clusters composed only of ERW samples (**Figure 1C and 1D**). In particular, the majority of ERW vOTUs assigned to the RNA virus families *Narnaviridae* (98.0%), *Botourmiaviridae* (84.2%), and *Cryppavirales* (56.4%) within the *Lenarviricota* phylum were found in ERW-specific clusters, suggesting that the diversity of soil viruses within these three families may be largely under-characterized (**Figure 2B**). Another 36.4% (for DNA phages) and 41% (for RNA viruses) were only clustered with other metagenome-derived virus genomes sampled from soil- and/or plant-related samples, and only 11 DNA and 220 RNA vOTUs were clustered with viruses from RefSeq database. Overall, these results indicate that the ERW phages and viruses are mostly novel compared to isolated references, but display some similarity to other uncultivated soil viruses, consistent with the existence of a “global soil virosphere” still only partially sampled^28^. This was confirmed for RNA viruses via UNIFRAC analyses of the different phylum-wide phylogenies, which indicated that ERW sequences overall were more closely related to sequences from other soil samples rather than sequences from other environments or references (**Figure 2B** and **2C, and Supplementary Table 11**).

Applying a new integrated phage-host prediction method (iPHoP^51^) which relies on an ensemble of phage-based and host-based approaches, 1,529 (37.8%) DNA vOTUs could be associated with a host genus or family (**Figure 1C and Supplementary Table 3**). Most of the predicted hosts were assigned to *Actinomycetia* (n = 618, 15.3%), *Alphaproteobacteria* (n = 506, 12.5%), and *Gammaproteobacteria* (n = 92, 2.27%), representing ∼80% of the total predicted hosts. On the other hand, the same approach (iPHoP) did not yield reliable host predictions for most RNA viruses, as this tool was primarily designed for DNA bacteriophages and archaeoviruses. Instead, we leveraged the RdRP phylogenies (see above) to identify putative hosts for RNA viruses, especially between bacteria and major divisions of eukaryotes. Overall, 2,449 (48.7%) RdRP branched within the *Leviviricetes* class and were assigned to prokaryote hosts, along with the 9 RdRP branching within the *Cystoviridae* family (**Figure 1D and Supplementary Table 4**). 723 RdRP branched within clades associated with fungal hosts (14.4%), including 31 (59.6%) and 41 (36.0%) clades in the *Duplornaviricota* and *Negarnaviricota* phyla, respectively (**Figure 1D, Figure 2B and Supplementary Figure 1A to 1D**). Finally, the rest of the RdRP were found in clades associated with various eukaryotic hosts (30%), or without any isolate representative (6.8%), highlighting the vast extent of soil RNA virus diversity still to be characterized.

### Contrasted dynamics of DNA and RNA viruses across seasons

We next investigated the dynamics of both DNA and RNA viral communities across seasons, depths and locations, using both presence/absence and nMDS ordination analyses based on estimated relative abundances (RPKM). Overall, 2,758 (68.2%) and 1,238 (24.6%) DNA and RNA viruses, respectively, were found at least in one sample of each season (**Figure 3A**). Conversely, 373 (9.2%) and 1,208 (24%) DNA and RNA viruses, respectively, were only found in a specific season (**Supplementary Figure 2A and 2B)**, indicating that both communities may exhibit seasonal dynamics, although of different magnitude. These patterns were confirmed by nMDS ordination analyses showing that both RNA (PERMANOVA; R^2^ = 0.19; *p < 0.001*) and DNA (PERMANOVA; R^2^ = 0.17; *p < 0.01*) viral community differed significantly by season (**Figure 3B and 3C**), while depth had only a marginal effect on the DNA viral community (PERMANOVA; R^2^ = 0.11; *p < 0.01*) and soil location had no significant effect on both communities (PERMANOVA; *p > 0.05*) (**Supplementary Table 12**). Bray-Curtis dissimilarities between months across successive seasons were also systematically significantly higher for the RNA viral community than for the DNA one, suggesting that RNA viruses underwent a higher rate of turnover between seasons (ANOVA; *p < 0.001*) (**Supplementary Figure 2C**). Finally, given the relative “stability” of the DNA virus community observed in metagenomes, we performed the same analysis for DNA vOTUs based on RPKM from metatranscriptomes, reasoning that transcriptional activity rather than relative abundance from metagenomes may uncover stronger seasonal patterns. Indeed, an nMDS ordination based on DNA vOTUs metatranscriptome RPKM was strongly structured by season (PERMANOVA; R^2^ = 0.34; *p < 0.001*) (**Figure 3D**). Altogether, these results indicate that both DNA and RNA virus communities are dynamic throughout the year, reflected primarily by a strong turnover for RNA viruses and changes in which subset of the community is transcriptionally active for the dsDNA phages.

**Figure 3.**
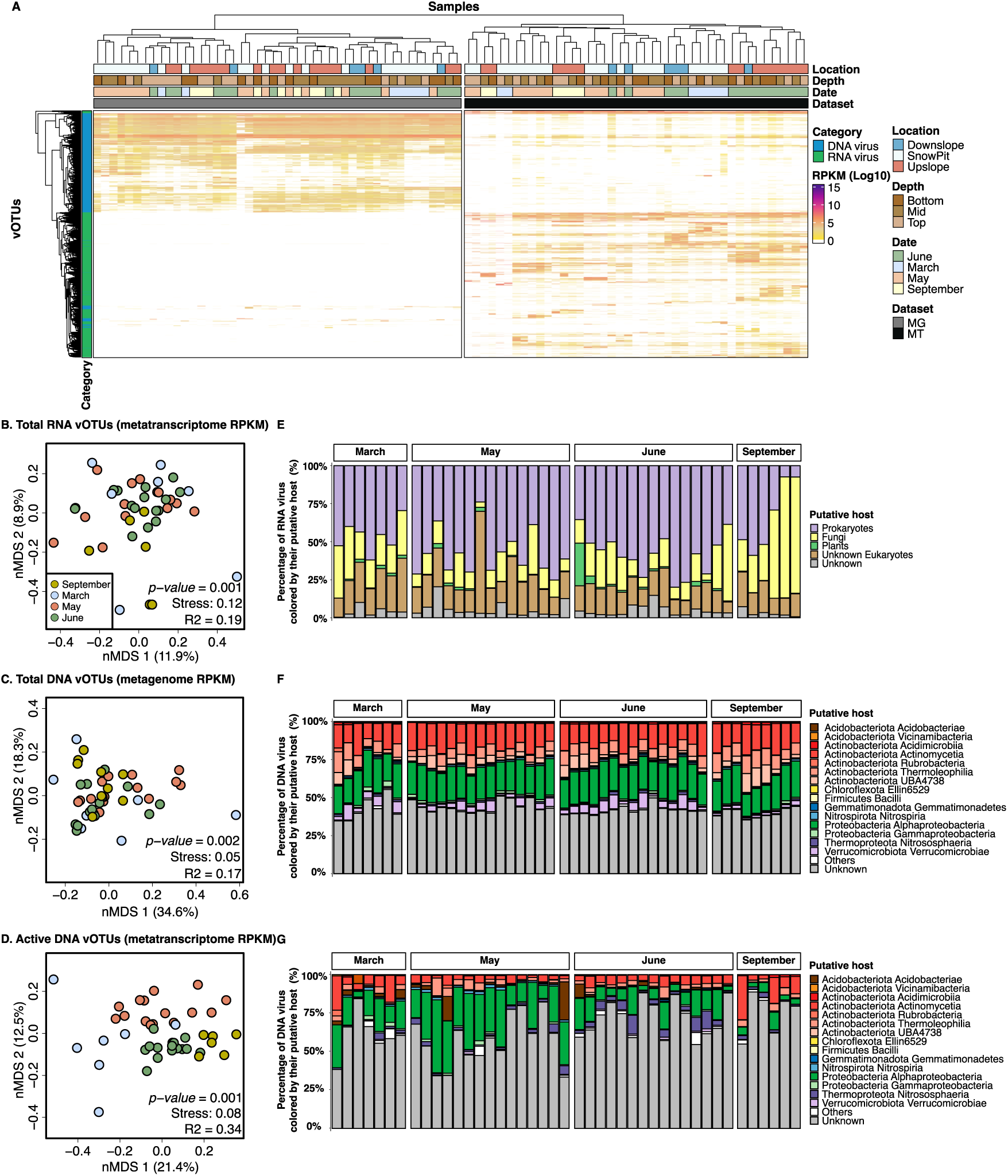
Overview of the temporal dynamics of total and active DNA and RNA viral communities. **A.** Distribution of all DNA (blue) and RNA (green) vOTUs across metagenome and metatranscriptome datasets. The vOTUs and samples are clustered based on vOTU relative abundances (log-transformed RPKM). Color bars above the heatmap indicate the location, depth, season, and type of each dataset. The left bar (STAMP) indicates the vOTUs with significant differences in relative abundance between seasons (*p* < 0.05). **B to G.** Beta-diversity of total RNA (B), total DNA (C) and active DNA (D) viral communities across the 4 seasons. For each group of viruses, non-metric multidimensional scaling (NMDS) ordination plots, representing the (dis)similarities between samples based on vOTU relative abundance, are presented in the left panels (B, C, and D). Individual samples are colored based on season: September (Yellow), March (Blue), May (Red), and June (Green). Stress values associated with two-dimensional ordination and PERMANOVA results describing the variance in community composition explained by season are reported for each plot. The relative abundance of RPKM of total RNA (E), total DNA (F) and active DNA (G) vOTUs predicted to infect putative host groups (for RNA vOTUs) or bacterial class (for DNA vOTUs) is indicated for each. “Other” represents remaining host classes (representing less than 0.1% of hosts). For “active” DNA, only vOTUs identified as active were considered (see Methods), and the RPKM from metatranscriptome read mapping was used as estimation of the relative abundance instead of RPKM from metagenome read mapping.

Some of these seasonal dynamics can be further explained by grouping vOTUs based on their predicted hosts. Throughout the year, the RNA virus community was characterized by an increased relative abundance of phages (mainly *Leviviricetes; Timlovirales*) in May during the snowmelt period, followed by an increase in relative abundance for plant-infecting viruses in June when the perennial plants emerge from dormancy, and for fungi-infecting viruses (*Cryppavirales*) in September (**Figure 3E and Supplementary Table 13**) following plant senescence and litter accumulation. When grouping DNA phages only a low proportion of phages predicted to infect *Thermoleophilia*, *Chloroflexota Ellin6529*, *Vicinamibacteria*, and *Verrucomicrobiae* bacterial hosts displayed a significant seasonal turnover (ANOVA; *p < 0.001*) (**Figure 3F and Supplementary Table 13**). Meanwhile, other groups of DNA phages found in both metagenome and metatranscriptome samples showed a significant seasonal dynamic based on their metatranscriptome RPKM, with a higher proportion of phages predicted to infect *Alphaproteobacteria, Acidimicrobiia, Verrucomicrobiae, Nitrospiria,* and *Actinomycetia* showing the highest change in activity over season (ANOVA; *p < 0.001*) (**Figure 3G and Supplementary Table 13**). These suggested that different types of virus-host interactions and ecological successions coexist in ERW soils. For example, broadly present and seasonally active viruses strongly influence the host community dynamics and structure. By contrast, seasonally dynamic but mostly inactive viruses do not affect host community structure, but are likely to be replicating alongside the host cell as part of lysogenic and/or chronic infections.

### Increased activity of DNA and RNA viruses after snowmelt

Because the viral populations were detected from bulk soil samples, they presumably represent a mixture of proviruses (i.e., integrated or extrachromosomal viruses that reside within their hosts cell), actively replicating viruses, and some extracellular viral particles^8^. Metatranscriptomes offer a unique opportunity to further understand which soil viruses are active, where, and when. To evaluate DNA viral activity at the vOTU level, metatranscriptomic libraries were used to identify expressed genes in each viral DNA genome, and viruses for which at least one expressed gene was detected were classified as active (**Figure 4A**). Among these, viral genomes that expressed genes related to virion structure, encapsidation, and/or lysis functions were classified as undergoing a lytic infection. The same approach is not possible for RNA viruses given their RNA-based genome, however for the dominant RNA viruses (the ssRNA *Lenarviricota* phylum), we instead leveraged strand-specific mapping information to identify actively replicating viruses based on the detection of both coding and non-coding genome strands (**Figure 4B**). This enabled a comparative analysis of activity levels for DNA and RNA viruses in ERW.

**Figure 4.**
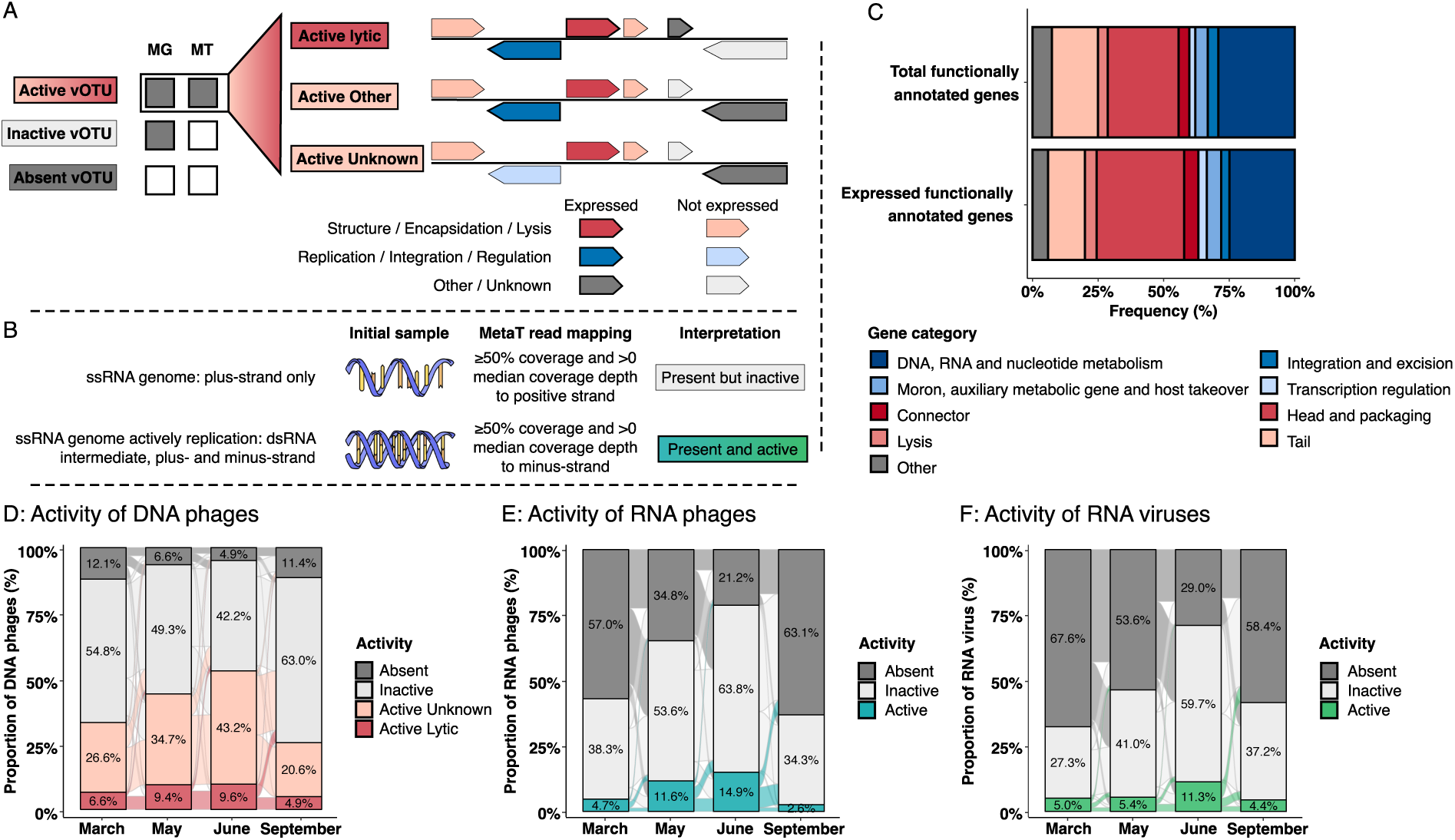
Functional annotation and activity of DNA and RNA viruses. **A.** Schematic representation of the framework used for assessing the activity, including infection stage, of DNA phages. Metatranscriptome read mapping is used to identify expressed genes in each viral DNA vOTU, and the number and annotation of these genes is then used to determine the activity and the infection stage of each DNA vOTU. **B.** Schematic representation of the framework used for assessing the activity of ssRNA viruses. Based on metatranscriptome read mapping, ssRNA viruses are classified as actively replicating if both coding and non-coding genome strands are detected, and considered as “present” if only the coding strand is detected. **C.** Proportion of total and expressed annotated genes based on functional annotation using Prokka v1.14.6 from DNA viral genomes by aligning them against PHROGS v4 database, with an e-value cut-off 1E-6. Functional categories associated with lytic infections, i.e., categories associated with virion production and host cell lysis, are colored in red, and the other major phage functional categories are colored in blue. Only genes that were annotated are included in the figure, and the proportion of annotated genes over all genes in the (active) DNA vOTUs is indicated next to each bar chart. **D.** Proportion of active (dark and light red), inactive (light grey), and absent (dark grey) DNA phages across months. Within DNA vOTUs identified as active, the ones likely engaged in active lytic infection was identified based on the functional annotation of expressed genes, while other active vOTUs are identified as “Active - Unknown”. **E and F.** Proportion of active (blue and green), inactive (light grey), and absent (dark grey) RNA phages (E) and RNA viruses (F) across months. A vOTU is considered as active when it is detected as active in at least one sample. The proportion of active vOTUs for each month is the sum of all active vOTUs for a given month.

Overall, 8,937 genes (31.5%) were identified as expressed in DNA viral genomes, including 1,101 functionally annotated (**Figure 4C and Supplementary Table 14**). A total of 2,926 (72.3%) DNA viral vOTUs were detected as active in at least one sample, and 535 (13.2%) were classified as active lytic viruses based on functional annotation (**Figure 4D and Supplementary Figure 3**). Meanwhile, for RNA viruses among the *Lenarviricota* phylum, 24.5% (600 vOTUs) of the *Leviviricetes* (**Figure 4E**) and 18.7% (294 vOTUs) of the eukaryote-infecting viruses (**Figure 4F**) were detected as active in at least one sample. Across seasons, the overall proportion of active DNA viruses significantly increased from September (25.6%) to June (52.8%). A similar overall pattern was recovered for both RNA bacteriophages (*Leviviricetes*) and other eukaryote-infecting *Lenarviricota*, with significant increases in activity from September (2.6% and 4.4%) to June (14.9% and 11.3%), suggesting that similar large-scale seasonal variations, here in particular the snowmelt followed by subsequent plant growth season, likely shape the activity of both DNA and RNA viral communities. Within this overall increase however, the highest increase in numbers of both active DNA and RNA phages occurred between March and May (peak snow to snowmelt period), while the largest increase in active RNA eukaryote-infecting viruses occurred between May and June (plant emergence post snowmelt), suggesting a delayed response to snowmelt for some eukaryotic viruses compared to phages.

### Active phages are driving the bacterial community structure in ERW soils

The increased activity observed for DNA phages throughout the year combined with the strong seasonal effect observed from metatranscriptome RPKM but not metagenome RPKM for the same DNA phages suggest that the ERW DNA phage community may be structured as a ‘seed-bank’. This ‘seed-bank’ is likely composed by a large group of persistent and mostly inactive phages residing in soils or in their host, and a different subset actively replicating and lysing their host across seasons, in particular following and/or concomitant with host growth/bloom.

To better characterize the relationship between host growth and viral activity, we first used similarity percentage breakdown (SIMPER)^52^ analyses to identify which DNA phage vOTUs were differentially active across seasons. Overall, 144 (5%) active DNA phages exhibited significant and clear differential abundance patterns in metatranscriptomes across seasons (ANOVA, Effect size > 0.3, adjusted *P* < 0.05), and could each be associated to a specific “high activity season” (**Figure 5A and Supplementary Table 15**). Further, 41.7% of these representative phages were assigned to a host taxon that was previously associated by Sorensen et al.^6^ to a specific ecological strategy (i.e., Winter-adapted, Snowmelt-specialist, and Spring-adapted taxa), which allowed us to explore the dynamics of active phage-host interactions across seasons (**Figure 5B and Supplementary Table 16**). In a somewhat counter-intuitive manner, DNA phages infecting both winter-adapted and snowmelt-specialist bacteria were least active when their predicted bacteria host would be growing, i.e., in March and May, respectively (**Figure 5A**). Conversely, DNA phages infecting spring-adapted bacteria were most active during the expected growth period of their hosts, between June and September. Phage-host interactions in the ERW thus appear to follow at least two distinct patterns: for the limited diversity of bacteria adapted to “slow” growth under snow or immediately upon snowmelt, the majority of DNA phage activity seems to be delayed compared to this initial seasonal growth of the host. On the other hand, the growth of a diverse community of bacteria in Spring following snowmelt seems to be associated with concomitant DNA phage activity. This suggests that the optimal infection strategy for soil bacteriophages may be, at least partially, driven by the ecological and growth strategy of their host. Finally, DNA phage activity may also respond directly to soil temperature variations across seasons, as soil temperatures may be near or above 0°C under snow cover and jump to 4°C in a short time period during the snowmelt period^6^.

**Figure 5.**
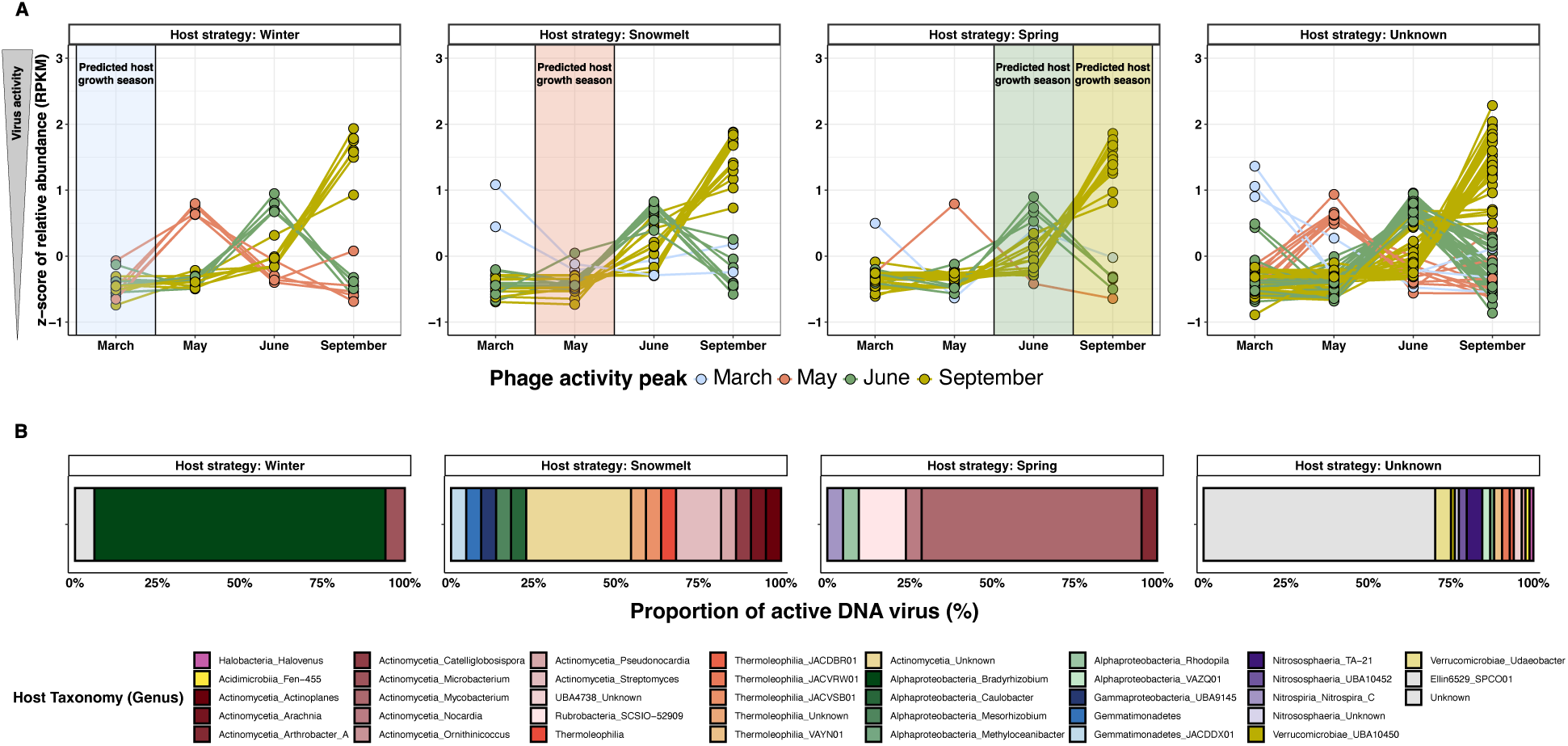
Temporal dynamics of virus-host relationships. **A.** Temporal dynamics of active DNA phages exhibiting a significant of changes in active DNA vOTU abundance between months (n = 144). The significance of changes in abundance between months was tested with a multiple group statistic test (ANOVA), a post-hoc test (Tukey-Kramer) to identify which pairs of months differ from each other and a multiple test correction (Storey’s FDR) to control false discovery rate, using STAMP. The seasonal dynamics of each active DNA vOTUs exhibiting significant changes in abundance between months was plotted using the mean of RPKM transformed in z-score. Each vOTU was associated to a specific season based on its peak of activity (colored lines). Finally, vOTUs’ dynamics are grouped by panel, depending of the “ecology strategy” of their assigned host (see Methods). Each host were associated to an “ecological strategy” depending to the month (or season) a given host was supposed to be growing^6^, represented by colored boxes in each panel. Finally, all active DNA vOTUs without assigned host or host without a clear ecological strategy were plotted in the last panel. **B.** Proportion of active DNA vOTUs exhibiting a significant of changes in active DNA vOTU abundance between months determined by STAMP, and colored by their putative host.

## DISCUSSION

Leveraging a data set of paired metagenomic and metatranscriptomic libraries from mountainous soil samples across seasons, we identified thousands of DNA and RNA viruses and assessed their diversity, community structure, and activity patterns. These data contribute to a better characterization of soil virus phylogenetic and genomic diversity, and suggest that soil virome composition and activity is affected by both their host metabolism and by ecological features of their local sampling environment. The ERW soils sampled in this study were characterized by a dominance of bacteriophages with two types of population: a large group of inactive phages residing in soils or in their hosts, and a smaller group of (often temporarily) active phages. The persistence of inactive viruses across seasons could have been facilitated by the continuous presence of putative hosts^32^, and/or low soil temperatures preventing viral inactivation^53^, especially under snow. Alternatively, it is possible that some of these persistent viruses maintain a low level of activity throughout the year not detectable in the current data. Regardless of the underlying mechanism of persistence, these phages likely represent a “seed-bank” from past lytic events that may serves as a reservoir for new infections to emerge when conditions become advantageous^54^. Given the abundance and diversity of hosts infected by these active viruses, the proliferation of these active and lytic phages will very likely have substantial impacts on the microbial communities, and thus on the soil biogeochemical cycling.

Phages are becoming increasingly recognized for their essential ecological roles, especially as they can control host population dynamics^26, 43, 55^. During lytic phases, virulent phages invade, replicate in and eventually kill their hosts, which can result in a substantial reduction in the relative abundances of the dominant microbial community members, as illustrated by the “Kill-The-Winner” (KtW) model^11, 12, 56–59^. In soils, temperate phages are considered to be more common, and have an opportunity to reside in their hosts via lysogenic infections rather than lysing them. This led to alternative virus-host dynamic models, such as for instance “Piggyback-The-Winner” (PtW), which predicts that phages integrate into their hosts’ genomes as prophages when microbial abundances and growth rates are high^55^. Finally, the factors determining the lytic to lysogenic switch in complex soil microbial communities remain poorly understood. In the ERW soils, lytic infection of active hosts seemed favored in spring and early summer (May / June), consistent with KtW interactions. This model is expected to occur under favorable conditions, such as a nutrient enrichment or wetting soils, generating high bacterial diversity^32^ and favoring lytic over lysogenic infections^60^, which is consistent with conditions during the plant-growing season in ERW soils. Conversely, the temporal delay between the activity of phages and the growing period of their hosts under snow conditions or upon snowmelt associated to the low diversity of hosts may be best described by a third model, recently proposed: “Piggyback-The-Loser” (PtL)^61^ or “Piggyback-The-Persistent” (PtP)^62^, that suggests the opposite of the PtW model. This hypothesis argues that when the host is growing slowly, it is more beneficial to remain in a lysogenic state, as it is less likely that the phages produced through lysis will find a new host^42^. Consistently, PtL or PtP was described in some regions of the deep sea or polar marine regions^61, 63^; and the dynamics observed in ERW during March and May suggest that comparable infection strategies also occur in soils. Furthermore, the sequential identification of different infection dynamics across seasons suggest that environmental conditions and host growth strategies may likely be an important factor driving the selection for one strategy or the other.

In terms of community diversity, while only few RNA bacteriophages have been experimentally characterized so far, we found a large number of RNA phages belonging to the *Leviviricetes* class in the ERW soils, including potentially novel phages associated with a wide range of bacteria inhabiting soils. In combination with recent investigations of *Leviviricetes* in terrestrial ecosystems, our study emphasizes the underappreciation of RNA phages in soils relative to DNA ones. This can be explained by the difficulties to recover RNA from soil because it is more easily degraded than DNA and/or because RNA viruses are not detected in metagenomes. We also identified a significant part of the RNA phage community as actively replicating, and given that all known RNA phages are virulent, we hypothesize that these significantly contribute to overall bacterial death in ERW^43^. By partially driving the turnover of bacteria, the diverse, abundant and active RNA phages recovered in this study are expected to impact soil microbiomes and associated biogeochemical cycling. Nevertheless, because of the limited availability of host predictions for RNA phages and the difficulties in linking them to even a high-order host taxon (e.g., a bacterial phylum or class), their impact on host communities in soil ecosystems remain difficult to evaluate more precisely. Further studies of global RNA phage diversity along with the development of new model systems, especially the cultivation of both RNA phages and their hosts, is now critical to understand the impact of these RNA phages on soil ecology^30, 64^.

Finally, beyond the apparent dominance of ssRNA *Leviviricetes* bacteriophages in ERW soils, our phylogenetic analysis suggests that a significant number of RNA viruses detected in this work infect fungal, plant and animal hosts. Among these, most of them display some similarity to other uncultivated soil viruses, with no match with isolated references, consistent with the existence of a “global soil virosphere” still only partially sampled^21, 29^. Interestingly, most of these viruses were predicted to infect fungal hosts, suggesting that the diversity of these mycoviruses may also be largely under-characterized^65^. Like the DNA and RNA phages, the eukaryotic-infecting RNA viruses seem to broadly follow their host dynamics, exhibiting a delayed peak of activity in comparison to phages. These results suggest that RNA viruses are an integral component of global soil ecosystems, with a diversity and activity driven by the growth of their hosts and the environmental parameters which affect them^66^. Altogether, our work demonstrates temporal viral activity dynamics, with time-shifted host-virus interactions depending of host strategies, viral resistance and coevolution^67^, highlighting the need to consider expanded models of host-virus interactions in further characterization and modeling of viral diversity, activity, and host impact in soil ecosystems.

## MATERIALS AND METHODS

### Field site description

The East River Watershed is located in Gunnison County, Colorado near the town of Crested Butte (38°57.5’ N, 106°59.3’ W) and is the location of the Rocky Mountain Biological Laboratory. Elevations within the East River watershed range from 2750 to 4000 m. Snow cover in winter typically persists 4 to 6 months (e.g., November through May) followed by an arid summer with intermittent, monsoonal rain events that occur from July through September. Minimum annual daily air temperature occurs in January (-14 ± 3° C), whereas the maximum daily air temperature typically occurs in July (23.5 ± 3° C). Plant composition at this location is a mixed, montane meadow community comprised of perennial bunchgrasses (e.g., *Festuca arizonica*), forbs (e.g., *Lupinus* spp., *Potentilla gracilis, Veratrum californicum*), and shrubs (*Artemesia tridentata*).

### Soil sampling

Soils were sampled from an upland hillslope location that was adjacent to the main stem of the East River (elevation ∼2775 m). Soil samples were collected using a 4 cm diameter soil bulk density corer on four dates starting at maximum winter snow depth (March 7, 2017), during the peak snowmelt period (May 9, 2017), during the plant growing season after the complete loss of snowpack (June 9, 2017), and lastly in autumn after plant senescence (September 16, 2017) (**Supplementary Table 1 and 2**). When there was no snowpack (September and June), soil samples were collected from 6 locations separated by approximately 10-meters. We dug three snow pits to the soil surface in March (approximately 1.5 meters of snowpack) and May (30 cm of snowpack) at locations adjacent to previous sampling sites. Two replicate soil cores separated by 1 meter were sampled from each snowpit as described above. All soil samples were split into three discrete depth increments; 0 to 5 cm, 5 to 15 cm, and 15 cm + below the soil surface. A ∼10 g subsample from each soil core at each depth was frozen immediately on dry ice in the field and archived at -80 °C for metagenome and metatranscriptome analyses (described below).

### Nucleic Acid Extraction

Nucleic acids were extracted on ice from 5 to 7 technical replicates of each soil sample by adding 0.5 mL phenol:chloroform:isoamyl alcohol (25:24:1) (Sigma-Aldrich, St. Louis, MO, USA) to 0.5 g of soil in a 2 ml Lysing Matrix E tube (MP Biomedicals, Solon, OH, USA), followed by addition of 0.5 mL of CTAB buffer (5% CTAB, 0.25M phosphate buffer pH 8.0, 0.3M NaCl) and 50 μL of 0.1M aluminum ammonium sulfate. The samples were homogenized at 5.5 m/s for 30 s in a FastPrep-24 instrument (MP Biomedicals, Solon, OH, USA), then centrifuged at 16K g for 5 min at 4°C. The aqueous phase was removed and transferred to MaxTract High Density 2 mL tubes (Qiagen Inc, Valencia, CA, USA). Samples were then extracted a second time as described above and the aqueous phase from the repeated extractions for each sample were combined. Sodium acetate (3M sodium acetate, 1/10^th^ volume of total aqueous phase) and ice-cold ethanol (100%, 2X volume of total aqueous phase) were added, the samples were vortexed briefly, and a crude nucleic acid pellet was precipitated overnight at -20 °C. Following overnight precipitation, technical replicates for each soil sample were combined, then separation of DNA and RNA was completed using the AllPrep DNA/RNA Mini Kit (Qiagen Inc, Valencia, CA, USA). The amount of DNA or RNA extracted was quantified using the Qubit 1X dsDNA Broad Range Kit or Qubit RNA High Sensitivity Kit, respectively (ThermoFisher Scientific). DNA and RNA quality were assessed using a 2100 Bioanalyzer instrument (Agilent Technologies, Santa Clara, USA). DNA and RNA was stored at -80 °C prior to metagenome and metatranscriptome analyses (described below).

### Library preparation and sequencing

Sequence data was generated at the DOE Joint Genome Institute (JGI) using Illumina technology. For metagenomes, library preparation for Illumina sequencing was performed using the Kapa Biosystems library preparation kit. Briefly, DNA was sheared using a Covaris LE220 focused-ultrasonicator. The sheared DNA fragments were size selected by double-SPRI and then the selected fragments were end-repaired, A-tailed, and ligated with Illumina compatible sequencing adaptors from IDT containing a unique molecular index barcode for each sample library. The prepared libraries were quantified using KAPA Biosystems’ next-generation sequencing library qPCR kit and run on a Roche LightCycler 480 real-time PCR instrument. Sequencing of the flowcell was performed on the Illumina NovaSeq sequencer using NovaSeq XP V1 reagent kits, S4 flowcell, following a 2x151 indexed run recipe. For metatranscriptomes, rRNA was depleted using the Illumina’s Ribo-Zero rRNA Removal Kit (Bacteria), and stranded cDNA libraries were generated using the Illumina TruSeq Stranded Total RNA kit. Briefly, the rRNA-depleted RNA was fragmented and reversed transcribed using random hexamers and SSII (Invitrogen) followed by second strand synthesis. The fragmented cDNA was treated with end-pair, A-tailing, adapter ligation, and 10 cycles of PCR. qPCR was used to determine the concentration of the libraries. The prepared libraries were quantified using KAPA Biosystems’ next-generation sequencing library qPCR kit and run on a Roche LightCycler 480 real-time PCR instrument. Sequencing of the flowcell was performed on the Illumina NovaSeq sequencer using NovaSeq XP V1 reagent kits, S4 flowcell, following a 2x151 indexed run recipe.

### Assembly and annotation of metagenomes and metatranscriptomes

Metagenome libraries were filtered and assembled using the DOE JGI Metagenome Workflow^68^. Briefly, BBDuk (version 38, https://jgi.doe.gov/data-and-tools/bbtools/) was used to remove contaminants via k-mer matching, trim reads that contained adapter sequence, right quality trim reads where quality drops to 0, and remove reads that contained 4 or more ’N’ bases or had an average quality score across the read less than 3 or had a minimum length <= 51 bp or 33% of the full read length. Reads aligned with BBMap (https://jgi.doe.gov/data-and-tools/bbtools/) to masked human, cat, dog and mouse references at 93% identity or to common microbial contaminants were excluded from downstream analyses. Specific commands and options used for read filtering are available for each library on the JGI Data portal (https://data.jgi.doe.gov). Filtered reads were error-corrected using bfc (version r181) with " bfc -1 -s 10g - k 21 -t 10^69^, and then assembled with metaSPAdes^70, 71^ version 3.13.0 using the “metagenome” flag, running the assembly module only (i.e., without error correction) with kmer sizes of 33, 55, 77, 99, and 127. Contigs larger or equal to 200 bp were then annotated using the IMG pipeline v.4.16.4^72^. Briefly, protein-coding genes were predicted with Prodigal v2.6.3^73^ and prokaryotic GeneMark.hmm v2.8^74^, and compared to COG 2003^75^, Pfam v30^76^, and IMG-NR 20180530^68^ using HMMER 3.1b2^77^ and lastal 914^78^ for taxonomic and functional annotation. Predicted protein-coding genes are also assigned to KEGG Orthology (KO) terms^79^ and Enzyme Commission (EC) numbers based on similarity to reference sequences in the IMG-NR 20180530 database^68^.

In addition to these individual assemblies of metagenome libraries, a combined assembly of the 48 metagenome libraries was performed using MetaHipmer^80^. Filtered reads from the 48 libraries were pooled and used as input for metagenome assembly with MetaHipmer v1, using default parameters. Contigs larger or equal to 500bp were then submitted to IMG for a similar functional and taxonomic annotation as previously described, using the IMG annotation pipeline v.5.0.23^68, 81^.

Metatranscriptome libraries were similarly processed using the default JGI workflow. BBDuk (version 38, https://jgi.doe.gov/data-and-tools/bbtools/) was used to remove contaminants and ribosomal RNA via k-mer matching, trim reads that contained adapter sequence, right quality trim reads where quality drops to 0, and remove reads that contained 1 or more ’N’ bases, had an average quality score across the read less than 10 or had a minimum length <= 51 bp or 33% of the full read length. Reads aligned with BBMap (https://jgi.doe.gov/data-and-tools/bbtools/) to masked human, cat, dog and mouse references at 93% identity, to common microbial contaminants, or to ribosomal RNA sequences were excluded from downstream analyses. Filtered reads were then de novo assembled with MEGAHIT v1.1.2^82^, using k-mer sizes of 23, 43, 63, 83, 103, and 123. Contigs larger or equal to 200bp were then submitted to IMG for annotation as previously described, using the IMG annotation pipeline v4.16.5^72^.

### Viral sequence detection and annotation

All contigs from the individual assemblies of metagenomes and metatranscriptomes and from the combined assembly of metagenomes were processed with VirSorter2 (v2.0.beta) for virus sequence detection^45^. These predictions were then further refined by identifying and removing potential host contaminants using CheckV v0.8.1^46^, inspecting all predicted proviruses, i.e. cases in which a contig is predicted to harbor both host and viral region(s) to refine provirus boundaries, and removing if necessary all predicted viral sequences with similarity to Type VI Secretion Systems. CheckV v0.8.1^46^ was then used to estimate the completeness of all filtered predicted viral sequences, and only sequences larger or equal to 5kb or estimated to be at least 50% complete (AAI-based completeness) were retained. For metatranscriptomes, contigs detected as likely RNA viruses based on a custom identification of RdRP genes^30^ were added to the filtered dataset obtained from the VirSorter 2 analysis.

The full dataset of predicted viral sequences was clustered into vOTUs following standard guidelines^47^ at 95% ANI and 85% AF using Mummer 4.0.0b2^83^, and the longest sequence was selected as the representative for each vOTU. To complement the IMG functional annotation, predicted protein-coding genes from selected representatives were compared to proteins from RefSeq Virus r2016^84^ using Diamond 0.9.24^85^ (minimum score of 50), to the VOGdb v205 (https://vogdb.org) using HMMER 3.3.2^77^ (minimum score of 30), and to the Pdb70 v190918^86^, Pfam v32^87^, and SCOPe70 v1.75^88^ databases (database package downloaded in Feb. 2019 from the HH-Suite website) using Hhblits 3.1.0^89^ (minimum probability of 90). Selected representatives were then further refined to identify and remove sequences only encoding putative Insertion Sequences based on annotation keywords “Transposase”, “insertion sequence”, and “insertion element”, as well as sequences without any annotated gene (i.e., only composed of predicted cds without any significant hit to any database). This led to a final dataset of 9,321 vOTUs.

### Estimation of the relative abundances of vOTUs

To evaluate vOTU and individual gene coverage, reads from individual metagenome and metatranscriptome libraries were mapped to the vOTU representative sequences with BBMap v38 (default parameters, https://jgi.doe.gov/data-and-tools/bbtools/). For metatranscriptomes, samtools v1.13^90^ and bedtools v2.30.0^91^ were used to calculate strand-specific coverage for both full sequences and individual genes. All reads mapped to vOTUs were used to calculate the RPKM (Reads Per Kilobase per Million mapped reads) value of each vOTU after normalizing by the sequence depth (per million reads) and the length of the contig (in kilobase).

### Phylogenetic analyses, taxonomic assignment and host prediction

GeNomad v1.0.0beta (https://github.com/apcamargo/genomad) was used for taxonomic classification (default parameters) of each vOTU^48^. iPHoP v0.9beta^51^ was used to predict the host family of each vOTU using a minimum score of 75, default parameters otherwise, and with the prediction with the highest score selected for each vOTU.

To refine the taxonomic classification and the host prediction of RNA vOTUs, phylogenetic analysis was performed using RdRP sequences. Previously published multiple sequence alignments of RdRPs^49, 92^ were used to search for RdRP sequences using HMMER 3.3.2^77^ on ERW viral contigs, recovered as described above, and on contigs produced by previous studies^13, 24, 49, 92^. We supplemented the data set with RdRP sequences collected from NCBI RefSeq Virus database r2016^84^ and group II intron reverse transcriptases (RT), used as an outgroup. Acceptance criteria for the RdRP profiles searches were E-value ≤1e-12 and score ≥ 50. This analysis identified 28,916 non-redundant contigs encoding predicted proteins with significant amino acid sequence similarity to previously identified RdRP. The extracted RdRP sequences were broadly assigned to the five major branches of RdRPs based on the best hit to the RdRP profiles^49^, and clustered using MMseqs2 v14^93^ with sequence similarity threshold of 0.5 and a coverage threshold of 80%. The representative sequences of each cluster were aligned using MAFFT v7.505^94^ (default parameters). All alignments were trimmed using TrimAl v1.3^95^ with the -gappyout option, and used to reconstruct maximum likelihood trees using FastTree v2.3^96^, and rooted by RT sequences. When visualizing phylogenetic trees, all branches with support values lower than 50 were collapsed.

The procedure to establish the RNA vOTU features (taxonomic affiliation and host prediction) across the trees (i.e., clades) relies on the tree topology and leverages existing taxonomic and host prediction information from RefSeq Virus r2016: all sequences belonging to a monophyletic clade in which all reference sequences are affiliated to a single taxon *T* or connected to a single host H were also assigned to the same taxon *T* or host H. A custom *perl* script was used to apply this logic based on the reference taxonomy and the host information for each phylogeny.

### Ecological distribution analyses

For DNA phages, a network analysis using vContact v2.0^50^ with “−rel-mode Diamond”, “−vcs-mode ClusterONE”, and all other settings set to default was used to compare the ≥10kb DNA ERW virus genomes, prokaryotic virus genomes from NCBI RefSeq Virus database (v94) r2016^97^ and more than 12,000 viral genomes from the viral database PIGEON v1.0^29^. ERW viral genomes were assigned into viral clusters (VCs) when clustering was significant (*p* < 0.05) and classified as outliers when clustering was non-significant. All unclustered viral genomes were classified as singletons.

For RNA viruses, previously identified ERW RdRP sequences and RdRP sequences from a custom database containing more than 613,000 RNA virus genomes from environmental metatranscriptomic studies and Refseq prokaryotic virus genomes (see above) were clustered using MMseqs2 v14^93^ with sequence similarity threshold of 0.5 to identify clusters unique to ERW or shared with other datasets. To complement this gene-based clustering analysis, generalized unweighted UNIFRAC distances were calculated using *GUniFrac* package on R, with *α* = 0.5 (parameter controlling weight on abundant lineages) to evaluate the distance between datasets based on the sequences included in the RdRP phylogeny analyses described above.

### Viral genome annotation, activity and infection cycle prediction

Genes were predicted and annotated using Prokka v1.14.6^98^ from viral genomes by aligning them against the PHROGS v4 database, with an e-value cut-off of 1E-6. To evaluate DNA viral activity, metatranscriptomic libraries were used to identify expressed genes in each viral DNA genome. A gene was considered as expressed if the coding strand had ≥50% of its positions covered and >0 median coverage depth. DNA vOTUs for which at least one expressed gene was detected were classified as active. Based on the genome annotation described above, active DNA vOTUs that expressed genes related to virion structure, encapsidation, and/or lysis functions were classified as undergoing a lytic infection while the other active DNA genomes were considered as possibly lysogenic or chronic infections, and broadly classified as “Active - Other”. Additionally, for DNA vOTUs identified in known bacteriophage/archaeovirus taxa, BacPhlip v0.9.6^99^ was used to predict lifestyle (i.e. temperate or lytic) based on the vOTU representative sequences.

While the same approach of identifying expressed genes in metatranscriptomes is not possible for RNA viruses given their RNA-based genome, we still evaluated the activity of the dominant ssRNA *Lenarviricota* viruses using the strand-specific mapping information of metatranscriptomes. The rationale was that, while these genomes are expected to be single-stranded, their replication intermediary should be double-stranded. Hence, RNA vOTUs with ≥50% of the genome covered at 1x or more on the coding strand with a >0 median coverage depth was considered as detected, while RNA vOTUs for which both strands were covered for ≥ 50% at 1x or more were classified as detected and active.

### Statistical analyses

All statistical analysis and figures were performed in R (CRAN)^100^ and Rstudio using the *vegan*, *ggplot2* and *ComplexUpset* packages, and STAMP (Statistical Analysis of Metagenomic Profiles) software v 2.1.3^101^. Non-metric multidimensional scaling (nMDS) ordination and Hierarchical clustering analysis based on Bray-Curtis dissimilarity matrices, using the *vegan* package, was used to visualize sample comparisons. Bray-Curtis dissimilarity matrices were also generated for both DNA and RNA viral communities to visualize the similarity between and within months. Analyses of variance (ANOVA) and permutational multiple analysis of variance (PERMANOVA) tests were used to identify significant differences in viral community composition between dates, depths, and locations. Finally, we tested the significance of changes in active DNA vOTU abundance between months with a multiple group statistic test (ANOVA), a post-hoc test (Tukey-Kramer) to identify which pairs of months differ from each other, and a multiple test correction (Storey’s FDR) to control false discovery rate, using STAMP. Post-hoc plots generated by STAMP show the results of each significant test (corrected p-value < 0.01), and provide an effect size measure for each pair of months. Based on the STAMP analyses, we then plotted the temporal dynamics (z-score of RPKM) of each active DNA vOTUs exhibiting significant changes in abundance between months based on the “ecological strategy” of the assigned host, using the iPHoP analyses (see above). Based on Sorensen et al.^6^, each host were associated to an “ecological strategy” depending on the month (or season) a given host was most abundant. All active DNA vOTUs without assigned host or assigned to a host without a clear ecological strategy were plotted in a separated panel.

### Data availability

All available metagenomic and metatranscriptomic data, are available through the IMG/M portal (assemblies) and NCBI SRA (reads). IMG/M and SRA identifiers of all metagenomes and metatranscriptomes, along with detailed information for each sample, are available in Supplementary Table 1 and 2.

## DECLARATION

## Acknowledgements.

We wish to thank Jenny Reithel and Shannon Sprott at Rocky Mountain Biological Laboratory (RMBL) for their general assistance at the field site.

## Author contributions

PS, and EB designed the overall study, which is a contribution to the Watershed Function Scientific Focus Area lead by Hubbard S. Susan. SR and CC designed the virus-specific study. Data collection was performed by PS (primarily) and SW, UK, and EB. CC, and SR performed the bioinformatic analyses, CC, SR, and EE-F wrote the manuscript. All authors contributed to revising the manuscript, and approved the final manuscript.

## Funding information

A portion of this research was performed under the Facilities Integrating Collaborations for User Science (FICUS) program and used resources at the DOE Joint Genome Institute and the Environmental Molecular Sciences Laboratory (grid.436923.9), which are DOE Office of Science User Facilities. Both facilities are sponsored by the Biological and Environmental Research program and operated under Contract Nos. DE-AC02-05CH11231 (JGI, https://ror.org/04xm1d337) and DE-AC05-76RL01830 (EMSL, https://ror.org/04rc0xn13). This material is based upon work supported as part of the Watershed Function Scientific Focus Area funded by the U.S. Department of Energy, Office of Science, Office of Biological and Environmental Research under Award Number DE-AC02-05CH11231. Rocky Mountain Biological Laboratory (RMBL) lab spaces and equipment are funded in part by the RMBL Equipment (Understanding Genetic Mechanisms) Grant, DBI-1315705. This work was supported by the U.S. Department of Energy, Office of Science, Biological and Environmental Research, Early Career Research Program awarded under UC-DOE Prime Contract DE-AC02-05CH11231.

## Conflicting interests

The authors declare that they have no conflict of interest.

## Notes

### Competing Interest Statement

The authors have declared no competing interest.

