## Supplementary Figures for "Virus diversity and activity is driven by snowmelt and host dynamics in a high-altitude watershed soil ecosystem"

**Figure S1. Diversity and phylogenetic analyses of ERW RNA viral communities**. Rooted phylogenetic trees of RdRP sequences belonging to the *Pisuviricota* (Figure S1A), *Kitrinoviricota* (Figure S1B), *Duplornaviricota* (Figure S1C), and *Negarnaviricota* (Figure S1D) phyla. The trees are rooted using reverse transcriptases as an outgroup and visualized with *ggtree.* Clusters composed exclusively of ERW sequences are colored in brown (ring 1) with branches leading to these clusters highlighted in light brown in the tree, while clusters composed of ERW sequences and existing virus sequences are colored by the environment type of the study (soil: dark brown, aquatic: blue, public databases: dark grey). Virus taxonomy (ring 3) and host (ring 4) are predicted based on the position of reference sequences from the RefSeq database in the tree (see Methods). Relative abundance of RPKM of *Pisuviricota* (Figure S1E), *Kitrinoviricota* (Figure S1F), *Duplornaviricota* (Figure S1G), and *Negarnaviricota* (Figure S1H) phyla.

A. *Pisuviricota*

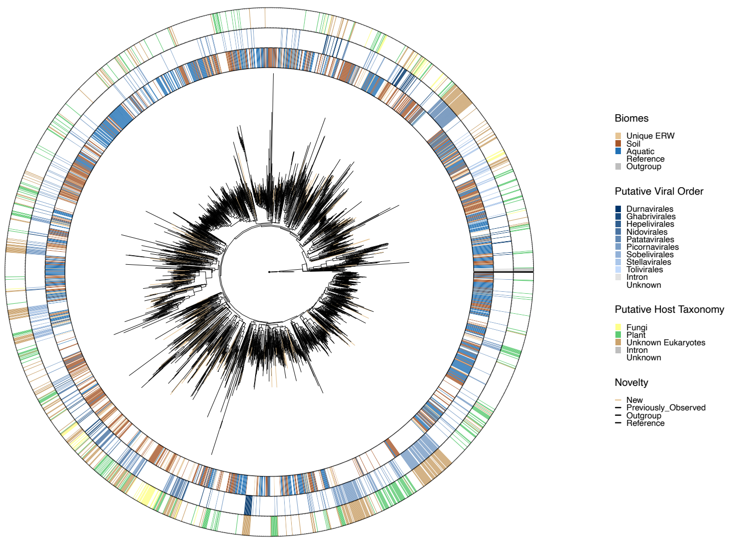


E

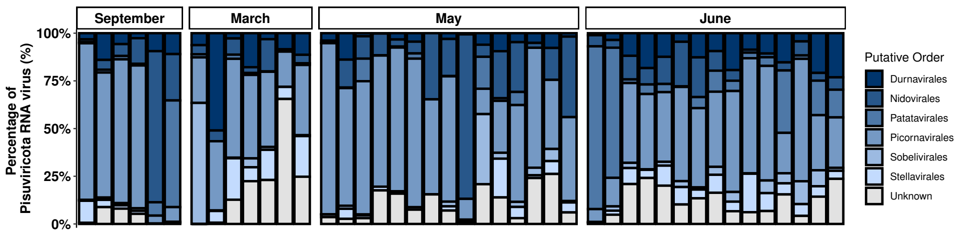


B. *Kitrinoviricota*

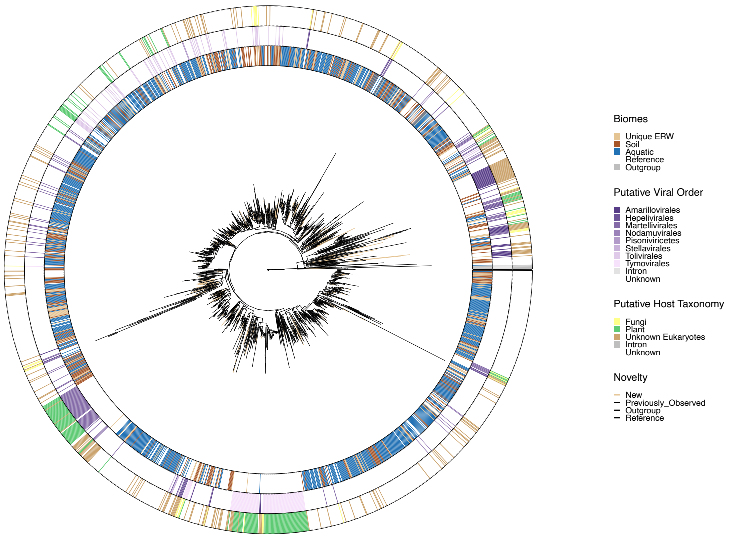


F

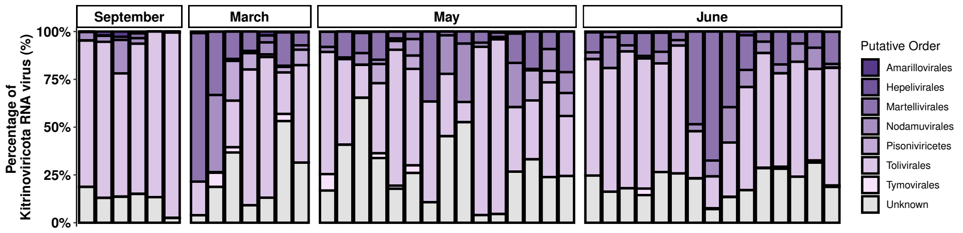

### C. *Duplornaviricota*

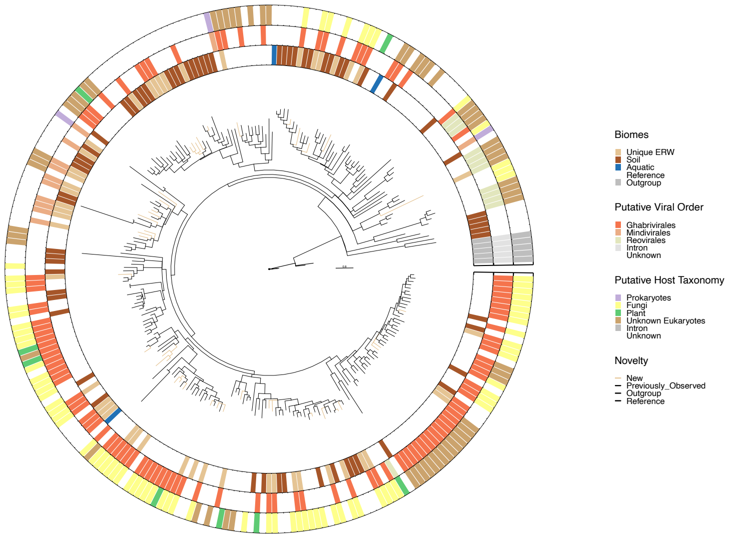


## G

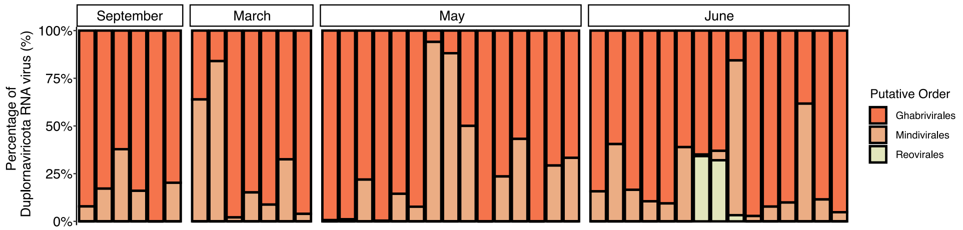


### D. *Negarnaviricota*

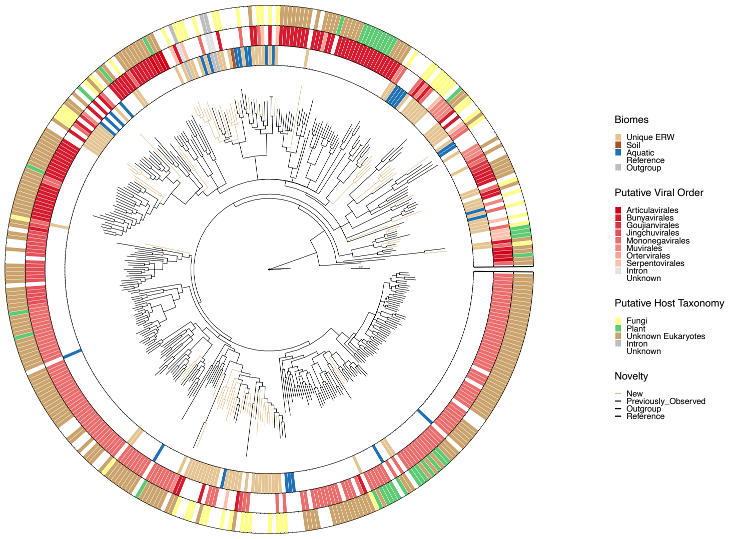


## H

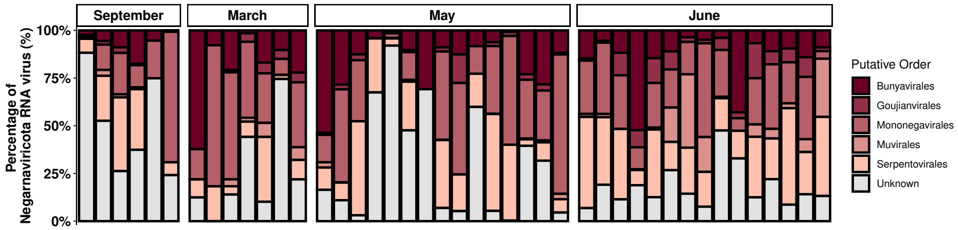

**Figure S2. Temporal dynamics of total and active DNA and RNA viral communities.** Upset plots of the distribution of DNA (Figure S2A) and RNA (Figure S2B) vOTUs by month, in which vOTUs are grouped based on the (combination of) month(s) they were detected in. Boxplots (Figure S2C) of Bray-Curtis dissimilarities between months and within months for DNA and RNA phages, and eukaryotes-infecting RNA viruses.

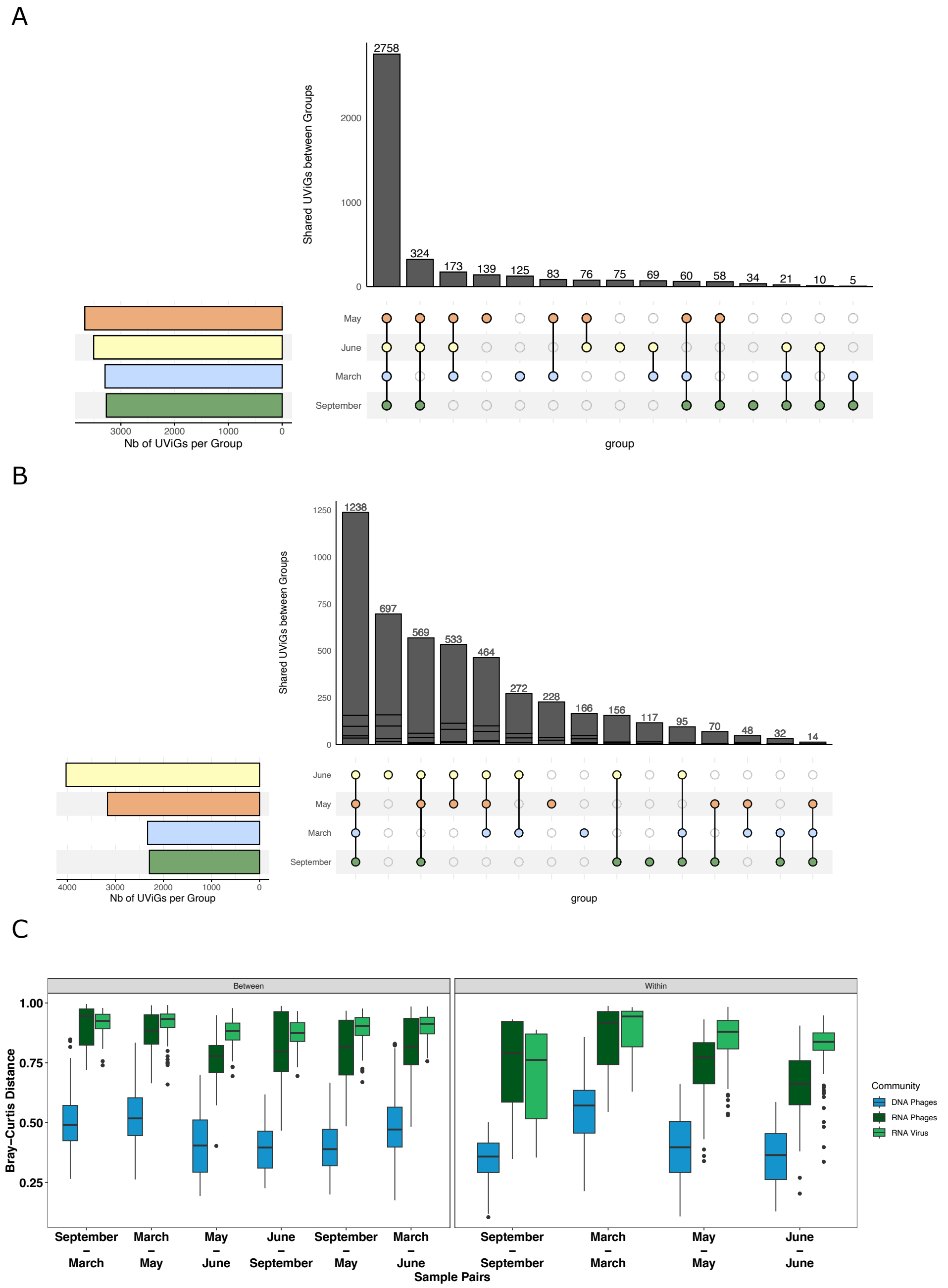

**Figure S3. Activity of DNA phages.** Proportion of active (dark and light red), inactive (light grey), and absent (dark grey) DNA phages across months for predicted temperate (left panel) and predicted lytic (right panel) DNA vOTUs using BACPHLIP. Within DNA vOTUs identified as active, the ones likely engaged in active lytic infection was identified based on the functional annotation of expressed genes, while other active vOTUs are identified as "Active - Unknown". A vOTU is considered as active in a given month when it is detected as active in at least one sample. The proportion of active vOTUs for each month is the sum of all active vOTUs for a given month.

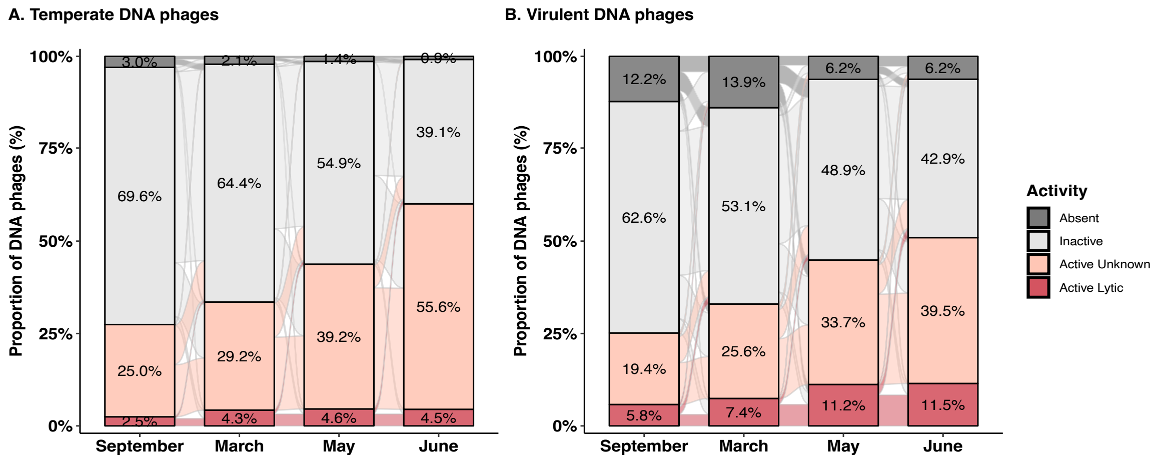
